# Structural basis of Gabija anti-phage defense and viral immune evasion

**DOI:** 10.1101/2023.05.01.538945

**Authors:** Sadie P. Antine, Alex G. Johnson, Sarah E. Mooney, Azita Leavitt, Megan L. Mayer, Erez Yirmiya, Gil Amitai, Rotem Sorek, Philip J. Kranzusch

## Abstract

Bacteria encode hundreds of diverse defense systems that protect from viral infection and inhibit phage propagation**^1–5^**. Gabija is one of the most prevalent anti-phage defense systems, occurring in >15% of all sequenced bacterial and archaeal genomes**^1,6,7^**, but the molecular basis of how Gabija defends cells from viral infection remains poorly understood. Here we use X-ray crystallography and cryo-EM to define how Gabija proteins assemble into an ∼500 kDa supramolecular complex that degrades phage DNA. Gabija protein A (GajA) is a DNA endonuclease that tetramerizes to form the core of the anti-phage defense complex. Two sets of Gabija protein B (GajB) dimers dock at opposite sides of the complex and create a 4:4 GajAB assembly that is essential for phage resistance *in vivo*. We show that a phage-encoded protein Gabija anti-defense 1 (Gad1) directly binds the Gabija GajAB complex and inactivates defense. A cryo-EM structure of the virally inhibited state reveals that Gad1 forms an octameric web that encases the GajAB complex and inhibits DNA recognition and cleavage. Our results reveal the structural basis of assembly of the Gabija anti-phage defense complex and define a unique mechanism of viral immune evasion.

---

Bacterial Gabija defense operons encode the proteins GajA and GajB that together protect cells against diverse phages^1^. To define the structural basis of Gabija anti-phage defense, we co- expressed *Bacillus cereus* VD045 GajA and GajB and determined a 3.0 Å X-ray crystal structure of the protein complex (Fig. 1a,b, Extended Data Fig. 1a,b, and Extended Data Table 1). The structure of the GajAB complex reveals an intricate 4:4 assembly with a tetrameric core of GajA subunits braced on either end by dimers of GajB (Fig. 1b). Focusing first on individual Gabjia protein subunits, GajA contains an N-terminal ATPase domain that is divided into two halves by insertion of a protein dimerization interface (discussed further below) (Fig. 1c). The GajA ATPase domain consists of an eleven-stranded β-sheet β1^ABC^, 2^ABC^, 4–6^ABC^ and β3^ABC^, 7–11^ABC^ that folds around the central α1^ABC^ helix (Fig. 1c, Extended Data Fig. 2). Sequence analysis of diverse GajA homologs demonstrates that the GajA ATPase domain contains a highly conserved ATP-binding site shared with canonical ABC ATPase proteins (Extended Data Fig. 2)^8^. The GajA C-terminus contains a four-stranded parallel β-sheet β1–4^T^ surrounded by three α-helices α3^T^, α4^T^, and α12^T^ that form a Toprim (topoisomerase-primase) domain associated with proteins that catalyze double-stranded DNA breaks (Fig. 1c, Extended Data Fig. 2)^9,10^. Consistent with a role in dsDNA cleavage, the structure of GajA confirms previous predictions of overall shared homology between GajA and a class of DNA endonucleases named OLD (overcoming lysogenization defect) nucleases^11,12^. Discovered initially as an *E. coli* phage P2 protein responsible for cell toxicity in *recB*, *recC* mutant cells^13–15^, OLD nucleases occur in diverse bacterial genomes as either single proteins (Class 1) or associated with partner UvrD/PcrA/Rep-like helicase proteins (Class 2), but the specific function of most OLD nuclease proteins is unknown^11,12^. GajA is a Class 2 OLD nuclease with the Toprim domain containing a complete active site composed of DxD after β3^T^ (D432 and D434), an invariant glutamate following β2^T^ (E379), and an invariant glycine between α1^T^ and β1^T^ (G409) similar to the active site of *Burkholderia pseudomallei* (*Bp*OLD) previously demonstrated to be essential for a two-metal-dependent mechanism of DNA cleavage (Fig. 1d, Extended Data Fig. 2)^11^.

**Figure 1.**
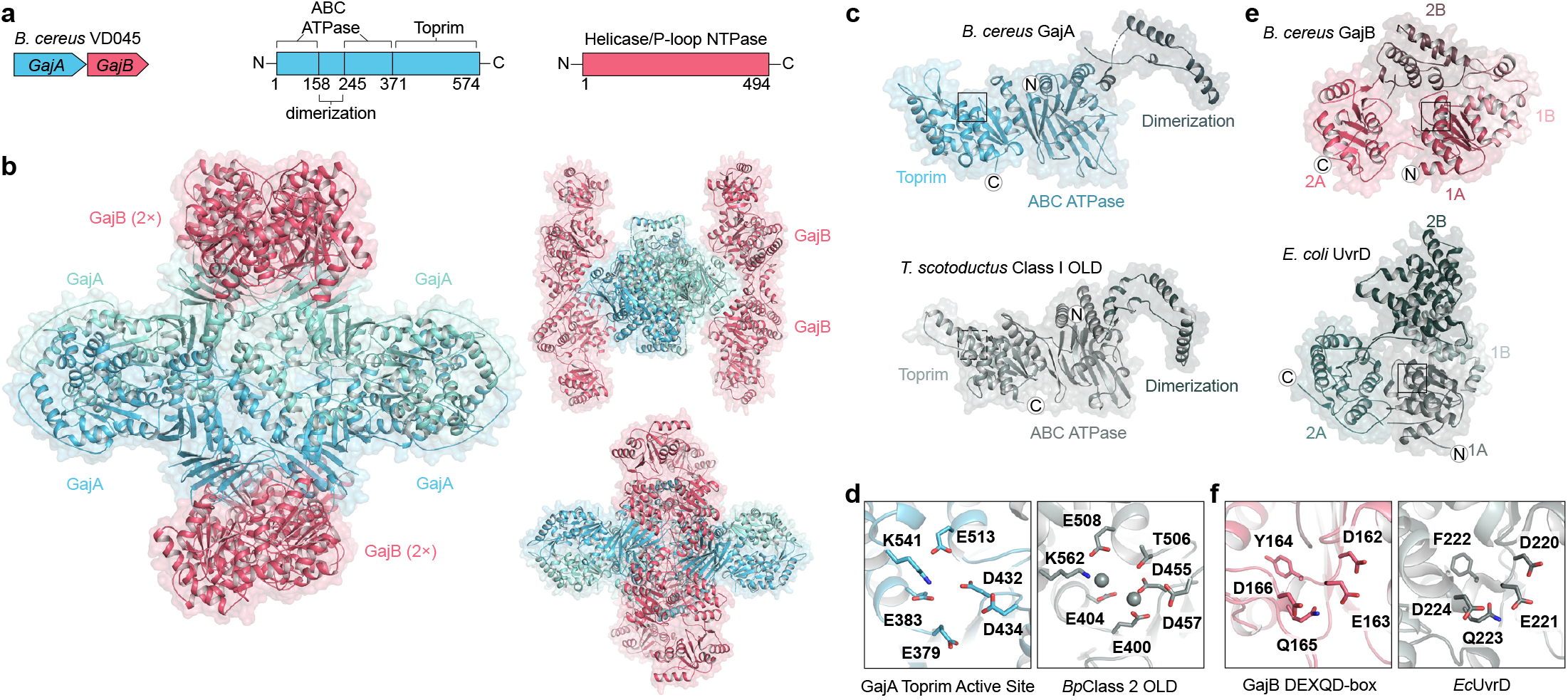
Structure of the Gabija anti-phage defense complex. **a,** Schematic of *B. cereus* Gabija defense operon and domain organization of GajA and GajB. **b,** Overview of the GajAB X-ray crystal structure shown in three orientations. GajA protomers are depicted in two shades of blue and GajB protomers are in red. **c,** Isolated GajA monomer (top) and comparison with a *Ts*OLD nuclease monomer (bottom) (Protein Data Bank (PDB) ID 6P74)^12^. **d,** Close-up view of GajA (left) and *Bp*OLD (right) (PDB ID 6NK8)^11^ Toprim catalytic residues. Location of GajA cutaway image is indicated with a box in (c) and magnesium ions are depicted as grey spheres. **e,** Isolated GajB monomer (top) and comparison with *Ec*UvrD (bottom) (PDB ID 2IS2)^20^. **f,** Close-up view of GajB (left) and *Ec*UvrD (right) DEXQD-box motif. Location of GajB cutaway image is indicated with a box in (e).

**Figure 2.**
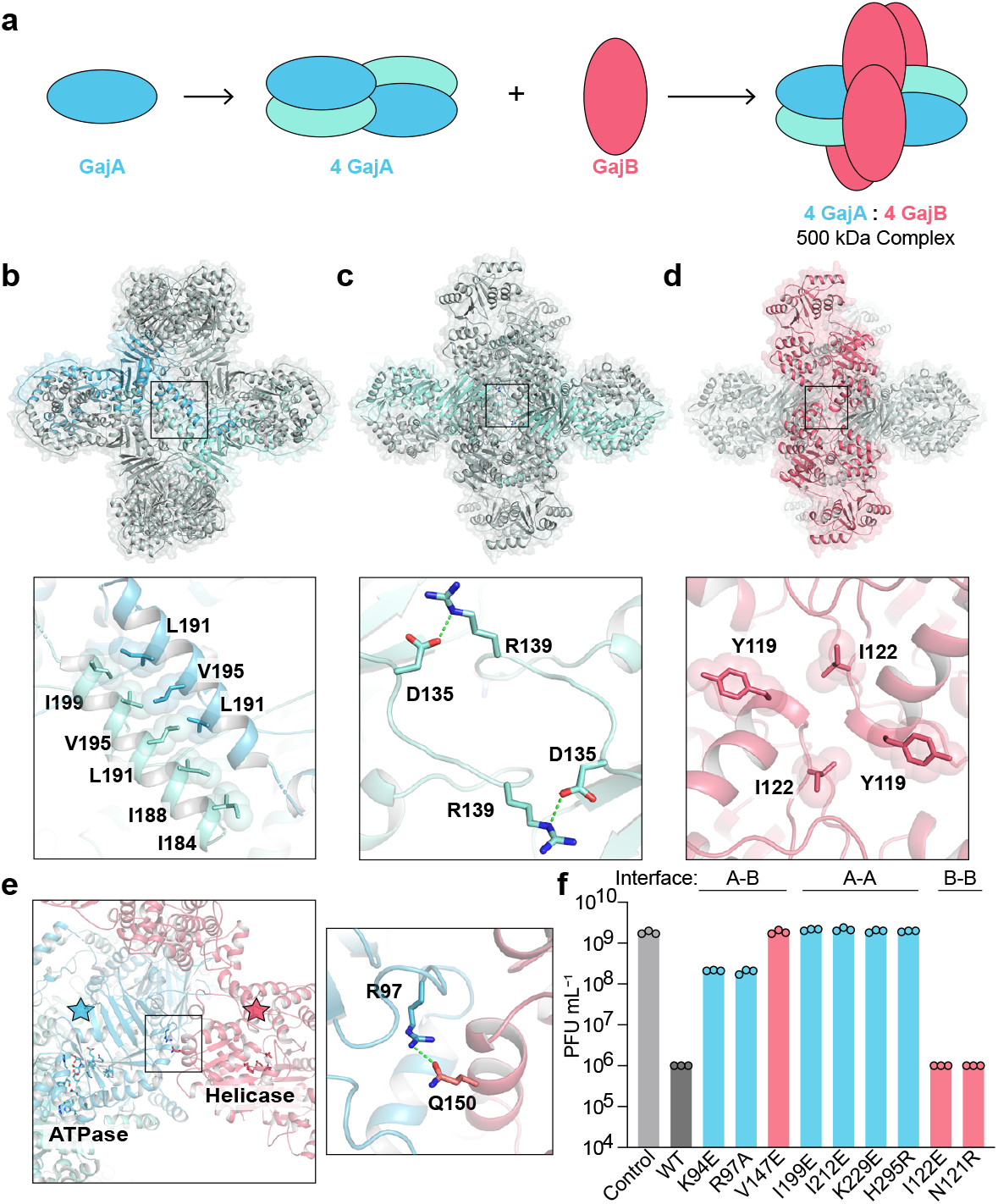
Mechanism of Gabija supramolecular complex assembly. **a,** Schematic model of GajAB complex formation by GajA tetramerization and GajB docking. **b,** Overview of the GajA α2^D^–α2^D^ dimerization interface and detailed view of interacting residues. For clarity, each GajA monomer is depicted in two shades of blue. **c,** Overview of the GajA–GajA ATPase interaction and detailed view of inter-subunit D135–R139 interaction. **d,** Overview of the minimal GajB–GajB dimer interface and detailed view of GajB–GajB hydrophobic interactions centered around Y119 and I122. **e,** Overview of the GajA–GajB interface highlighting proximity of GajA ABC ATPase and GajB helicase active site residues (left) with box indicating location of GajA R97 and GajB Q150 interaction (right). **f,** Analysis of GajA and GajB mutations in the GajA– GajB (A–B), GajA–GajA (A–A), and GajB–GajB (B–B) multimerization interfaces on the ability of the *B. cereus* Gabija operon to defend cells against phage infection. Data represent the phage SPβ average plaque-forming units (PFU) mL^−1^ of three biological replicates with individual data points shown.

The structure of GajB reveals a Superfamily 1A DNA helicase domain typically occurring in bacterial DNA repair (Fig. 1a,b)^16^. Superfamily 1A helicase proteins like UvrD, Rep, and PcrA translocate along ssDNA in the 3′–5′ direction, and are architecturally divided into four subdomains 1A, 1B, 2A, and 2B that reposition relative to each other during helicase function^16^. GajB contains all conserved helicase motifs required for ATP hydrolysis and nucleic acid unwinding including a Walker A motif Gx(4)GK-[TT] and a UvrD-like DEXQD-box Walker B motif responsible for NTP hydrolysis (Fig. 1f and Extended Data Fig. 3a)^16–18^. Activation of Superfamily 1A DNA helicase proteins like UvrD and Rep is known to require protein dimerization and rotation of the 2B subdomain^19–21^. Comparisons with UvrD and Rep demonstrate that GajB protomers in the GajAB complex exhibit partial rotation of the 2B domain relative to 2A-1A-1B consistent with a partially active conformation poised to interact with phage DNA (Fig. 1e and Extended Data Figure 1d).

**Figure 3.**
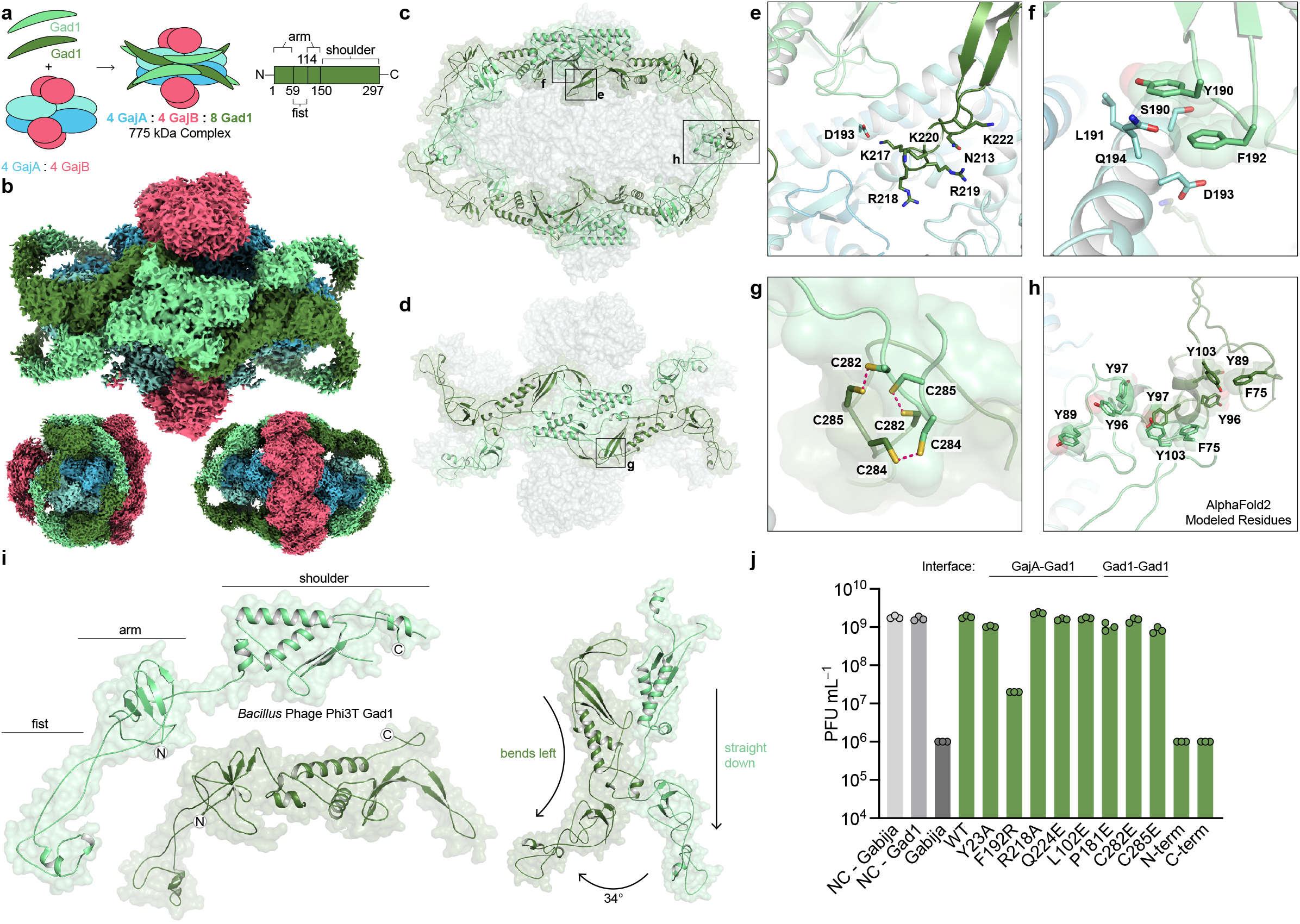
Structural basis of viral evasion of Gabija defense. **a,** Schematic model of GajAB–Gad1 co-complex formation and domain organization of phage Phi3T Gad1. **b,** Cryo-EM density map of *Bc*GajAB in complex with Phi3T Gad1 shown in three different orientations. The map is colored by the model, with Gad1 monomers depicted in two shades of green. **c,** Side-view of the complete Gad1 octameric complex and **d,** top-down view of the Gad1 tetrameric interface with boxes highlighting close-up views in (e–h). **e,f,** Zoomed-in views of major Gad1–GajA interface contacts including a Gad1 positively charged loop (e) and hydrophobic interactions with GajA α2^D^ (f). **g,h,** Zoomed-in views of major Gad1–Gad1 oligomerization interactions including disulfide bonds in the C-terminal shoulder domain (g) and fist–fist domain contacts modeled by rigid-body placement of an AlphaFold2 fist domain structure prediction into the cryo-EM map (h). **i,** Two distinct conformations of Gad1 observed in the GajAB– Gad1 co-complex structure. Differences in Gad1 arm domain rotation are highlighted on the right. **j,** Analysis of Gad1 mutations in the GajA–Gad1 and Gad1–Gad1 multimerization interfaces on the ability Gad1 to enable evasion of Gabija defense. Data represent PFU mL^−1^ of phage SPβ infecting cells expressing *Bc*Gabija and *Shewanella sp.* phage 1/4 Gad1, or negative control (NC) cells expressing either plasmid empty vector. *Shewanella sp.* phage 1/4 Gad1 residues are numbered according to the Phi3T Gad1 structure. Data are the average of three biological replicates with individual data points shown.

## Mechanism and function of Gabija supramolecular complex formation

To define the mechanism of Gabija complex assembly, we analyzed oligomerization interfaces within the GajAB structure. Purification of individual Gabija proteins demonstrates that GajA is alone sufficient to oligomerize into a homo-tetrameric assembly (Extended Data Fig. 1b). GajB migrates as a monomer on size-exclusion chromatography, supporting a stepwise model of GajAB assembly (Fig. 2a and Extended Data Fig. 1b). GajA tetramers form through two highly-conserved oligomerization interfaces (Fig. 2b,c and Extended Data Fig. 2). First, the GajA N-terminal ATPase domain contains an insertion between β7^ABC^ and β8^ABC^ that consists of four α- helices (α1–4^D^) that zip-up against a partnering GajA protomer to form a hydrophobic interface along the α2^D^ helix (Fig. 2b). A similar α1–4^D^ dimerization interface exists in the structure of the bacterial *T. scotoductus* Class 1 OLD (*Ts*OLD) protein demonstrating that this interface is conserved within divergent OLD nucleases (Figs. 1c and 2c)^12^. The GajA ATPase domain contains a second oligomerization interface in a loop between β6^ABC^ and α6^ABC^ where hydrogen bond contacts between D135 and R139 interlock two GajA dimers to form the tetrameric core assembly (Fig. 2c). Compared to GajA, the GajB–GajB dimerization interface is minimal and consists of a hydrophobic surface in the GajB helicase 1B domain centered at Y119 and I122 (Fig. 2d). Major GajA–GajB contacts also occur in the GajB helicase 1B domain where GajA R97 in a loop between α4^ABC^ and β5^ABC^ forms hydrogen-bond contacts with Q150 in GajB α7 along with hydrophobic packing interactions centered at GajB V147 (Fig. 2d and Extended Data Fig. 3a). Notably, the GajAB structure demonstrates that the GajB helicase 1A subdomain including the DEXQD-box active-site is positioned adjacent to the GajA ATPase domain suggesting that GajB ATP-hydrolysis and DNA unwinding activity may regulate GajA ATPase domain activation (Fig. 2e). In addition to the major GajAB interface contacts, Gabija supramolecular complex assembly is driven by extensive protomer interactions that result in ∼31,000 Å^2^ of surface area buried for the GajA tetramer and ∼1,800 Å^2^ of surface area buried for each GajB subunit.

We reconstituted Gabija activity *in vitro* and observed that the GajAB complex rapidly cleaves a previously characterized 56 bp dsDNA substrate containing a sequence specific motif derived from phage lambda DNA (Extended Data Fig. 1c)^22^. GajA and GajB proteins are each essential for phage defense *in vivo*^1,22^, but we observed *in vitro* that GajA alone is sufficient to cleave target DNA suggesting a specific role for the GajAB complex in substrate recognition or nuclease activation during phage infection (Extended Data Fig. 1c). To confirm these findings, we tested a panel of GajAB interface mutations and measured the impact of substitutions on the ability of Gabija to defend *B. subtilis* cells from phage SPβ infection. Substitutions to the GajA– GajA dimerization interface including I199E, I212E, and K229E resulted in complete loss of phage resistance (Fig. 2f). Likewise, substitutions to the GajA–GajB hetero-oligomerization interface including GajA K94E, R97A and GajB V147E dramatically reduced the ability of Gabija to inhibit phage replication *in vivo*. In contrast, phage resistance was tolerant to mutations in the GajB– GajB interface suggesting that this minimal interaction surface is not strictly essential for anti-phage defense. Together, these results define the structural basis of GajA and GajB interaction and demonstrate that GajAB supramolecular complex formation is critical for Gabija anti-phage defense.

## Structural basis of viral inhibition of Gabija anti-phage defense

To overcome host immunity, phages encode evasion proteins that specifically inactivate anti-phage defense^23–28^. Yirmiya, Leavitt, and colleagues report discovery of the first viral inhibitor of Gabija anti-phage defense (Yirmiya and Leavitt et al 2023 Submitted Manuscript), and we reasoned that defining the mechanism of immune evasion would provide further insight into Gabija complex function. Gabija anti-defense 1 (Gad1) is a *Bacillus* phage Phi3T protein that is atypically large (35 kDa) compared to other characterized phage immune evasion proteins (Extended Data Fig. 4). Protein interaction analysis demonstrated that Gad1 binds directly to GajAB (Extended Data Fig. 5a,b), and we used cryo-EM to determine a 2.7 Å structure of the GajAB–Gad1 co-complex assembly (Fig. 3a,b, Extended Data Figs. 6 and 7a–f, and Extended Data Table 2). The GajAB–Gad1 co-complex structure reveals a striking mechanism of inhibition where Gad1 proteins form an oligomeric web that wraps 360° around the host defense complex. Eight copies of phage Gad1 encircle the GajAB assembly, forming a 4:4:8 GajAB–Gad1 complex that is ∼775 kDa in size (Fig. 3b,c). Gad1 primarily recognizes the GajA nuclease core, forming extensive contacts along the surface of the GajA dimerization domain (Fig. 3c,d). Key GajAB– Gad1 contacts include hydrogen-bond interactions from a Gad1 positively-charged loop located between β6 and β7 with GajA α2^D^ (Fig. 3e and Extended Data Fig. 8) and hydrophobic packing interactions between Gad1 Y190 and F192 with GajA α2^D^ (Fig. 3f and Extended Data Fig. 8). Although Gad1 contacts with GajB are limited, both GajA and GajB proteins are necessary for Gad1 interaction, demonstrating that Gad1 specifically targets the fully assembled GajAB complex to inactivate host anti-phage defense (Extended Data Fig. 5c).

**Figure 4.**
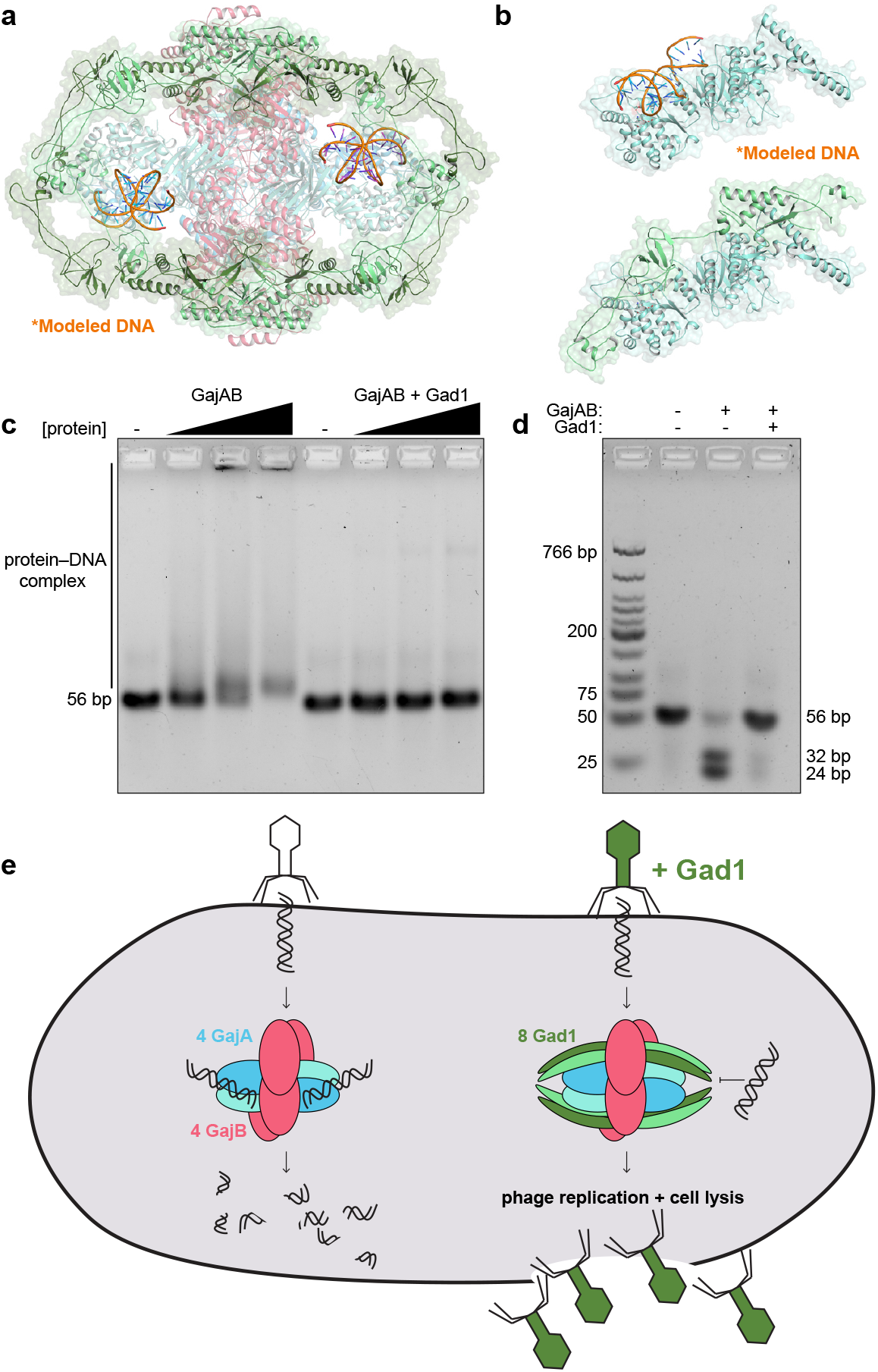
Inhibition of Gabija DNA binding and cleavage enables viral evasion. **a,** Cartoon representation of the GajAB–Gad1 co-complex structure with modeled DNA based on structural homology with *E. coli* MutS (PDB ID 3K0S)^31^. **b,** Isolated GajA protomer with modeled DNA bound to the Toprim domain (top) and same GajA promoter with Gad1 demonstrating significant steric clashes between Gad1 and the path of DNA (bottom). **c,d,** Biochemical analysis of GajAB 56-bp target DNA binding (c) and target cleavage (d) demonstrates that Gad1 potently inhibits GajAB activity. Data are representative of three independent experiments. **e,** Model of Gabija anti-phage defense and mechanism of Gad1 immune evasion.

Gad1 wraps around the GajAB complex using a network of homo-oligomeric interactions and remarkable conformational flexibility. On either side of the GajAB complex, four copies of Gad1 interlock into a tetrameric interface along the primary GajA binding site (Fig. 3d). The Gad1 tetrameric interface is formed by hydrogen-bond interactions between the C-termini “shoulder” domain of each Gad1 monomer and a highly conserved set of three cysteine residues C282, C284, and C285 that form disulfide interactions deep within an inter-subunit interface (Fig. 3d,g and Extended Data Fig. 8). The N-termini of each Gad1 monomer forms an “arm” domain that extends out from the shoulder and reaches around the GajA nuclease active site to connect to a partnering Gad1 protomer from the opposite side of the complex. At the end of the Gad1 arm is an N-terminal “fist” domain that allows two partnering Gad1 protomers to interact and complete the octameric web assembly (Fig. 3c,h). Particle heterogeneity limits resolution in this portion of the cryo-EM map, but AlphaFold2 modeling^29,30^ and rigid-body placement of the Gad1 N-terminal fist domain suggests conserved hydrophobic residues around the Gad1 α1 helix mediate the fist–fist interactions (Fig. 3h and Extended Data Fig. 8). To fully encircle GajAB, Gad1 adopts two distinct structural conformations. Each pair of Gad1 proteins that wrap around and connect at the GajAB complex edge are formed by one Gad1 protomer reaching out from the shoulder with an arm domain extended straight down and one Gad1 protomer reaching out with an arm domain bent ∼34° to the left (Fig. 3i and Extended Data Fig. 7h). Sequence analysis of Gad1 proteins from phylogenetically diverse phages demonstrates that the Gad1 N-terminal arm domain is highly variable in length (Extended Data Fig. 8), further supporting that conformational flexibility in this region is critical to inhibit host Gabija defense.

To test the importance of individual GajAB–Gad1 interfaces, we next analyzed a series of Gad1 substitution and truncation mutants for the ability to interact with GajAB and inhibit Gabija anti-phage defense. A Gad1 substitution F192R between β4 and β5 at the center of the primary GajA–Gad1 interface disrupted all ability of Gad1 to interact with GajAB *in vitro* and inhibit Gabjia anti-phage defense *in vivo* (Fig. 3j and Extended Data Fig. 9a). However, individual mutations throughout the periphery were insufficient to disrupt Gad1 inhibition of Gabjia anti-phage defense, demonstrating that the large footprint of Gad1 is tolerant to small perturbations that may enable host resistance. Likewise, mutations to the conserved Gad1 cysteine residues in the tetrameric shoulder interface greatly reduced stability of the GajAB–Gad1 complex formation *in vitro* but exhibited an ∼3-fold difference and still permitted Gad1 to block phage defense in *B. subtilis* cells (Fig. 3j and Extended Data Fig. 9a). Finally, in contrast to wildtype Gad1, expression of the Gad1 N-terminal fist–arm or C-terminal shoulder domains alone were unable to inhibit Gabija, demonstrating that full wrapping of Gad1 around the GajAB complex is necessary to enable phage evasion of anti-phage defense (Fig. 3j and Extended Data Fig. 9a).

## Inhibition of Gabija DNA binding and cleavage enables viral evasion

To define the mechanism of Gad1 inhibition of Gabija anti-phage defense, we next modeled interactions between GajAB and target DNA. The GajA Toprim domain is structurally homologous to the *E. coli* protein MutS involved in DNA repair^31^. Superimposing the MutS–DNA structure revealed positively charged patches lining a groove in the GajA Toprim domain that dips into the nuclease active site (Extended Data Fig. 10). Notably, the Gad1 arm domain directly occupies this putative DNA-binding surface supporting a model where the phage protein directly clashes with the path of target dsDNA (Fig. 4a,b). To determine the impact of viral inhibition on GajAB catalytic function we tested the role of Gad1 in individual steps of DNA binding and target DNA cleavage. Gad1 prevented GajAB from binding to target DNA and abolished all nuclease activity *in vitro* (Fig. 4c,d). Mutant Gad1 proteins F192R and C282E were no longer able to inhibit DNA cleavage, agreeing with the complete loss of F192R and reduced ability of C282E mutant proteins to block Gabija defense *in vivo* and form stable GajAB–Gad1 complexes *in vitro*. (Extended Data Fig. 9b). Together, these results demonstrate that phage Gad1 binds and wraps around the GajAB complex to block target DNA degradation and define a complete mechanism for immune evasion of Gabija anti-phage defense (Fig. 4e).

Our study defines the structural basis of Gabija supramolecular complex formation and explains how phages block DNA cleavage to defeat this form of host immunity. Similar to supramolecular complexes in CRISPR^32^, CBASS^33,34^, and RADAR immunity^35,36^, the ∼500 kDa GajAB complex extends an emerging theme in anti-phage defense where protein subunits assemble into large machines to resist phage infection. These results parallel human innate immunity, where key effectors in inflammasome, Toll-like receptor, RIG-I-like receptor, and cGAS-STING signaling pathways also oligomerize into large assemblies to block viral replication^37,38^. In contrast to the exceptionally large host defense complexes, phage evasion proteins are typically small 5–20 kDa proteins that sterically occlude key protein binding and active site motifs^24,25^. Breaking this rule, the 35 kDa anti-Gabija protein Gad1 is one of the largest described viral protein–protein inhibitors of host immune signaling (Extended Data Fig. 4). Whereas most viral evasion proteins >20 kDa in size are enzymatic domains that catalytically modify target host factors or signaling molecules, the large size of Gad1 is necessary to bind, oligomerize, and encircle around the entire host GajAB complex. Resistance to small phage proteins that simply block the GajA active site may explain why Gabija is a highly prevalent defense system in diverse bacterial phyla. Additionally, a key question opened by our structures of the Gabija complex is how GajB helicase activity is linked to activation of the GajA nuclease domain to control DNA target cleavage. Gad1 encasing the GajAB complex to trap it in an inactive state reveals a new mechanism for evasion of host defense and provides a key template to understand how viruses disrupt the complex mechanisms of activation of diverse anti-phage defense systems in bacteria.

## Methods

### Bacterial strains and phages

*B. subtilis* BEST7003 was grown in MMB (LB supplemented with 0.1 mM MnCl_2_ and 5 mM MgCl_2_) with or without 0.5% agar at 37°C or 30°C respectively. Whenever applicable, media were supplemented with ampicillin (100 μg mL^−1^), chloramphenicol (34 μg mL^−1^), or kanamycin (50 μg mL^−1^) to ensure the maintenance of plasmids. *B. subtilis* phages phi3T (BGSCID 1L1) and SPβ (BGSCID 1L5) were obtained from the Bacillus Genetic Stock Center (BGSC). Prophages were induced using Mitomycin C (Sigma, M0503).

Phage titer was determined using the small drop plaque assay method^39^. 400 µL of overnight culture of bacteria was mixed with 0.5% agar and 30 mL MMB and poured into a 10 cm^2^ plate followed by incubation for 1 h at room temperature. In cases of bacteria expressing Gad1 homolog and Gad1 mutations, 0.1–1mM IPTG was added to the medium. 10-fold serial dilutions in MMB were performed for each of the tested phages and 10 µL drops were put on the bacterial layer. After the drops had dried up, the plates were inverted and incubated at room temperature overnight. Plaque forming units (PFUs) were determined by counting the derived plaques after overnight incubation and lysate titer was determined by calculating PFU mL^−1^. When no individual plaques could not be identified, a faint lysis zone across the drop area was considered to be 10 plaques. Efficiency of plating (EOP) was measured by comparing plaque assay results on control bacteria and bacteria containing the defense system and/or a candidate anti-defense gene.

### Plasmid Construction

For protein purification and biochemistry, *B. cereus* VD045 *GajA* (IMG ID 2519684552) and *GajB* (IMG ID 2519684553) genes were codon-optimized for expression in *E. coli* and synthesized as gBlocks (Integrated DNA Technologies) and cloned into custom pET vectors with an N-terminal 6×His-SUMO2 fusion tag (GajB alone) or a C-terminal 6×His tag (GajA alone). GajA and GajB proteins were co-expressed together using custom pET vector with an N-terminal 6×His-SUMO2 or N-terminal 6×His-SUMO2-5×GS tag on GajA and ribosome binding site between GajA and GajB. Phi3T and *Shewanella sp.* phage 1/4 Gad1 (IMG ID 2708680195) gBlocks were cloned into a custom pBAD vector containing a chloramphenicol resistance gene and IPTG-inducible promoter. For Gad1 pull-down assays, *Shewanella sp.* phage 1/4 Gad1 was cloned with a ribosome binding site after the GajB gene in the N-terminal 6×His-SUMO2-5×GS GajAB plasmid. For plaque assays, the DNA of Gad1 was amplified from phage phi3T genome using KAPA HiFi HotStart ReadyMix (Roche cat # KK2601). Since Gad1 was toxic in *B. subtilis* cells containing Gabija, *Shewanella sp.* phage 1/4 Gad1 was used and synthesized by Genscript. Gad1 and related homologs were cloned into the pSG-thrC-Phspank vector^40^ and transformed to DH5α competent cells. The cloned vector and the vector containing Gad1 substitution and truncation mutants were subsequently transformed into *B. subtilis* BEST7003 cells containing Gabija integrated into the amyE locus^1^, resulting in cultures expressing both Gabija and a Gad1 homolog. As a negative control, a transformant with an identical plasmid containing GFP instead of the anti-defense gene, was used. Transformation in *B. subtilis* was performed using MC medium as previously described^1^. Sanger sequencing was then applied to verify the integrity of the inserts and the mutations. The pSG1 plasmids containing point mutations in Gabija were constructed by restriction-enzyme subcloning Gabija sequence into pGEM9Z, site-directed mutagenesis as previously described^41^, Gibson back into pSG1, and transformed into *B. subtilis* BEST7003 cells. Sanger sequencing of the mutations regions was then applied to verify the mutations in Gabija.

### Protein expression and purification

Recombinant GajAB and GajAB–Gad1 complexes were purified from *E. coli* as previously described^42^. Briefly, expression plasmids described above were transformed into BL21(DE3) or BL21(DE3)-RIL cells (Agilent), plated on MDG media plates (1.5% Bacto agar, 0.5% glucose, 25 mM Na_2_HPO_4_, 25 mM KH_2_PO_4_, 50 mM NH_4_Cl, 5 mM Na_2_SO_4_, 0.25% aspartic acid, 2–50 μM trace metals, 100 μg mL^−1^ ampicillin, 34 μg mL^−1^ chloramphenicol) and grown overnight at 37°C. Five colonies were used to inoculate 30 mL of MDG starter overnight cultures (37°C 230 rpm). 10 mL of MDG starter cultures were then inoculated in 1 L M9ZB expression cultures (47.8 mM Na_2_HPO_4_, 22 mM KH_2_PO_4_, 18.7 mM NH_4_Cl, 85.6 mM NaCl, 1% Cas-Amino acids, 0.5% glycerol, 2 mM MgSO_4_, 2–50 μM trace metals, 100 μg mL^−1^ ampicillin, 34 μg mL^−1^ chloramphenicol) and induced with 0.5 mM IPTG after reaching an OD_600_ of ≥1.5 (overnight, 16°C, 230 rpm).

After overnight induction, cells were pelleted by centrifugation, resuspended, and lysed by sonication in 60 mL lysis buffer (20 mM HEPES pH 7.5, 400 mM NaCl, 10% glycerol, 20 mM Imidazole, 1 mM DTT). Lysate was clarified by centrifugation, and supernatant was poured over Ni-NTA resin (Qiagen). Resin was then washed with lysis buffer, lysis buffer supplemented to 1 M NaCl, lysis buffer again, and finally eluted with lysis buffer supplemented to 300 mM Imidazole. Samples were then dialyzed overnight in 14 kDa MWCO dialysis tubing (Ward’s Science) with SUMO2-cleavage by hSENP2 as previously described^29,30^. hSENP2 did not efficiently cleave N- terminal 6×His-SUMO2-GajAB and the complex was therefore purified with an additional 5×GS linker. Proteins for crystallography and cryo-EM were dialyzed in dialysis buffer (20 mM HEPES- KOH pH 7.5, 250 mM KCl, and 1 mM DTT), purified by size exclusion chromatography using a 16/600 Superdex 200 column (Cytiva) and stored in gel filtration buffer (20 mM HEPES-KOH pH 7.5, 20 mM KCl, and 1 mM TCEP-KOH). Proteins for biochemical assays were dialyzed in dialysis buffer, purified by size exclusion chromatography using a 16/600 Superdex 200 column (Cytiva) or 16/600 Sephacryl 300 column (Cytiva) and stored in gel filtration buffer with 10% glycerol. Purified proteins were concentrated to >10 mg mL^−1^ using a 30 kDa MWCO centrifugal filter (Millipore Sigma), aliquoted, flash frozen in liquid nitrogen, and stored at −80°C.

For Gad1 pull-down assays, SUMO2-5×GS-GajA-GajB-Gad1 point mutant plasmids were transformed and expressed in BL21(DE3)-RIL cells and subject to Ni-NTA column chromatography. Proteins were dialyzed overnight along with SUMO2 cleavage with SENP2. Gad1 pulldown was analyzed by SDS-PAGE and Coomassie Blue staining.

### Crystallization and X-ray structure determination

Crystals were grown in hanging drop format using EasyXtal 15-well trays (NeXtal). Native GajAB crystals were grown at 18°C in 2 μL drops mixed 1:1 with purified protein (10 mg mL^−1^, 20 mM HEPES 250 mM KCl, and 1 mM TCEP-KOH) and reservoir solution (100 mM HEPES-NaOH pH 7.5, 2.4% PEG-400, and 2.2 M ammonium sulfate). Crystals were grown for 7 days before cryo- protection with reservoir solution supplemented with 25% glycerol and harvested by plunging in liquid nitrogen. X-ray diffraction data were collected at the Advanced Photon Source (beamlines 24-ID-C and 24-ID-E). Data were processed using the SSRL autoxds script (A. Gonzalez, Stanford SSRL). Experimental phase information was determined by molecular replacement using monomeric GajA and GajB AlphaFold2 predicted structures^29,30^ in Phenix^43^. Model building was completed in Coot^22^ and then refined in Phenix. The final structure was refined to stereochemistry statistics as reported in Extended Data Table 1. Structure images and figures were prepared in PyMOL.

### Electrophoretic mobility shift assay

56-bp sequence-specific motif dsDNA (5′ TTTTTTTTTT TTTTTTTAAT AACCCGGTTA TTTTTTTTTT TTTTTTTTTT TTTTTT 3′)^22^ was incubated with a final concentration of 2, 5, or 10 µM purified GajAB or GajAB–Gad1 complexes in 20 µL gel shift reactions containing 1 µM dsDNA, 5 mM CaCl_2_, and 20 mM Tris-HCl pH 8.0 for 30 min at 4°C. 10 µL was then mixed with 2 μL of 50% glycerol and separated on a 2% TB (Tris-borate) agarose gel. The gel was then run at 250 V for 45 min, post-stained with TB containing 10 µg mL^−1^ ethidium bromide rocking at room temperature, de-stained in TB buffer for 40 min, and imaged on ChemiDoc MP Imaging System.

### DNA cleavage assay

The same 56-bp dsDNA as above was incubated with GajAB or GajAB–Gad1 complexes in a 20 μL DNA cleavage reaction buffer containing 1 µM dsDNA, 1 µM GajAB or GajAB–Gad1, 1 mM MgCl_2_, 20 mM Tris-HCl pH 9.0 for 20 min at 37°C. Following incubation, reactions were stopped with DNA loading buffer containing EDTA and 10 µL was analyzed on a 2% TB agarose gel, which was run at 250V for 45 min. The gel was then post-stained rocking at room temperature with TB buffer containing 10 µg mL^−1^ ethidium bromide, de-stained in TB buffer alone for 40 min, and imaged on a ChemiDoc MP Imaging System.

### Cryo-EM sample preparation and data collection

For the SUMO2-GajAB–Gad1 co-complex sample, 3 μL of 1 mg mL^−1^ was vitrified using a Mark IV Vitrobot (Thermofisher). Prior to sample vitrification, 2/1 Carbon Quantfoil^TM^ grids were glow discharged using an easiGlow^TM^ (Pelco). Grids were then double-sided blotted for 9s, constant force of 0, 100% relative humidity chamber at 4°C, and a 10 s wait time prior to liquid ethane plunge and storage in liquid nitrogen.

GajAB–Gad1 co-complex cryo-EM grids were screened using a Talos Arctica microscope (Thermofisher) operating at 200 kV and the final map was collected on a Titan Krios microscope (ThermoFisher) operating at 300 kV. Both microscopes operated with a K3 direct electron detector (Gatan). SerialEM software version 3.8.6 was used for all data collection. For final data collection a total of 9,243 movies were taken at a pixel size of 0.3115 Å, a total dose of 41.1 e− /Å^2^, dose per frame of 0.63 e− /Å^2^ at a defocus range of range of −0.8 to −1.9 µm.

### Cryo-EM data processing

SBGrid Consortium provided data-processing software. Movies were imported into cryoSPARC^45^ for patch-based motion correction, patch-based CTF estimation, 2D and 3D particle classification, and non-uniform refinement. cryoSPARC data processing is outlined in Extended Data Figure 6. Briefly, after patch-based CTF estimation, five hundred micrographs were selected and autopicked using Blob Picker, which resulted in 625,295 particles after extracting from micrographs. 2D classifications were then used to generate 5 templates for Template Picker from which 110,654 particles were picked from 500 micrographs. After three more rounds of 2D classification 648,298 particles from all 9,243 micrographs were used in ab initios (K = 3), followed by heterogenous refinement. The best class with 573,410 particles was then used to go back and extract from all micrographs, which resulted in 570,485 particles that were used in non-uniform refinement resulting in a 2.86 Å C1 symmetry and 2.73 Å C2 symmetry map, which was then used for model building.

### Cryo-EM model building

Model building was performed in Coot^44^ by manually docking AlphaFold2 predicted structures^29,30^ as starting models and then manually completing refinement and model correction. To model the Gad1 fist domain, an AlphaFold2 model of the Gad1 arm–fist region was superimposed on the cryo-EM density of the manually built shoulder–arm region and then fit into density in Coot^44^. To complete the model for the sparse GajB density, the X-ray GajB structure was superimposed on the cryo-EM density. GajAB–Gad1 model was refined in Phenix^43^, and the structure stereochemistry statistics are reported in Extended Data Table 2. Figures were prepared in PyMOL and UCSF ChimeraX^46^.

### Statistics and reproducibility

Experimental details regarding replicates are found within figure legends.

## Acknowledgements

The authors are grateful to J. Asnes, J. Grippen, and members of the Kranzusch lab and Sorek lab for helpful comments and discussion and A. Lu for assistance with X-ray data collection. The work was funded by grants to P.J.K. from the Pew Biomedical Scholars program, the Burroughs Wellcome Fund PATH program, The Mathers Foundation, The Mark Foundation for Cancer Research, the Cancer Research Institute, the Parker Institute for Cancer Immunotherapy, and the National Institutes of Health (1DP2GM146250-01) and grants to R.S. from the European Research Council (ERC-AdG GA 101018520), the Israel Science Foundation (MAPATS Grant 2720/22), the Ernest and Bonnie Beutler Research Program of Excellence in Genomic Medicine, the Deutsche Forschungsgemeinschaft (SPP 2330, grant 464312965), and the Knell Family Center for Microbiology. E.Y. is partially supported by the Israeli Council for Higher Education (CHE) via the Weizmann Data Science Research Center. A.G.J. is supported through a Life Science Research Foundation postdoctoral fellowship of the Open Philanthropy Project. X-ray data were collected at the Northeastern Collaborative Access Team beamlines 24- ID-C and 24-ID-E (P30 GM124165), and used a Pilatus detector (S10RR029205), an Eiger detector (S10OD021527) and the Argonne National Laboratory Advanced Photon Source (DE- AC02-06CH11357). Cryo-EM data were collected at the Harvard Cryo-EM Center for Structural Biology at Harvard Medical School. We thank Theo Humphreys at PNCC for help with cryo-EM data collection. A portion of this research was supported by NIH grant U24GM129547 and performed at the PNCC at OHSU and accessed through EMSL (grid.436923.9), a DOE Office of Science User Facility sponsored by the Office of Biological and Environmental Research.

## Author Contributions

The study was designed and conceived by S.P.A. and P.J.K. All protein purification and biochemical assays were performed by S.P.A. and S.E.M. Crystallography structural analysis was performed by S.P.A. Cryo-EM structural analysis was performed by S.P.A., A.G.J., and M.L.M. Model building and analysis was performed by S.P.A. and P.J.K. Bioinformatics and protein sequence analysis was performed by E.Y., A.L., G.A, and R.S. Phage challenge assays were performed by A.L. and R.S. Figures were prepared by S.P.A. with assistance from S.E.M. The manuscript was written by S.P.A. and P.J.K. All authors contributed to editing the manuscript and support the conclusions.

## Additional Information

Correspondence and requests for materials should be addressed to P.J.K.

## Data Availability Statement

Coordinates and structure factors of the Gabija GajAB complex have been deposited in PDB under the accession code 8SM3. Coordinates and density maps of the GajAB–Gad1 co-complex are being deposited with PDB and EMDB under accession codes NNNN and NNNN. All other data are available in the manuscript or the supplementary materials.

## Extended Data Figure Legends

**Extended Data Figure 1.**
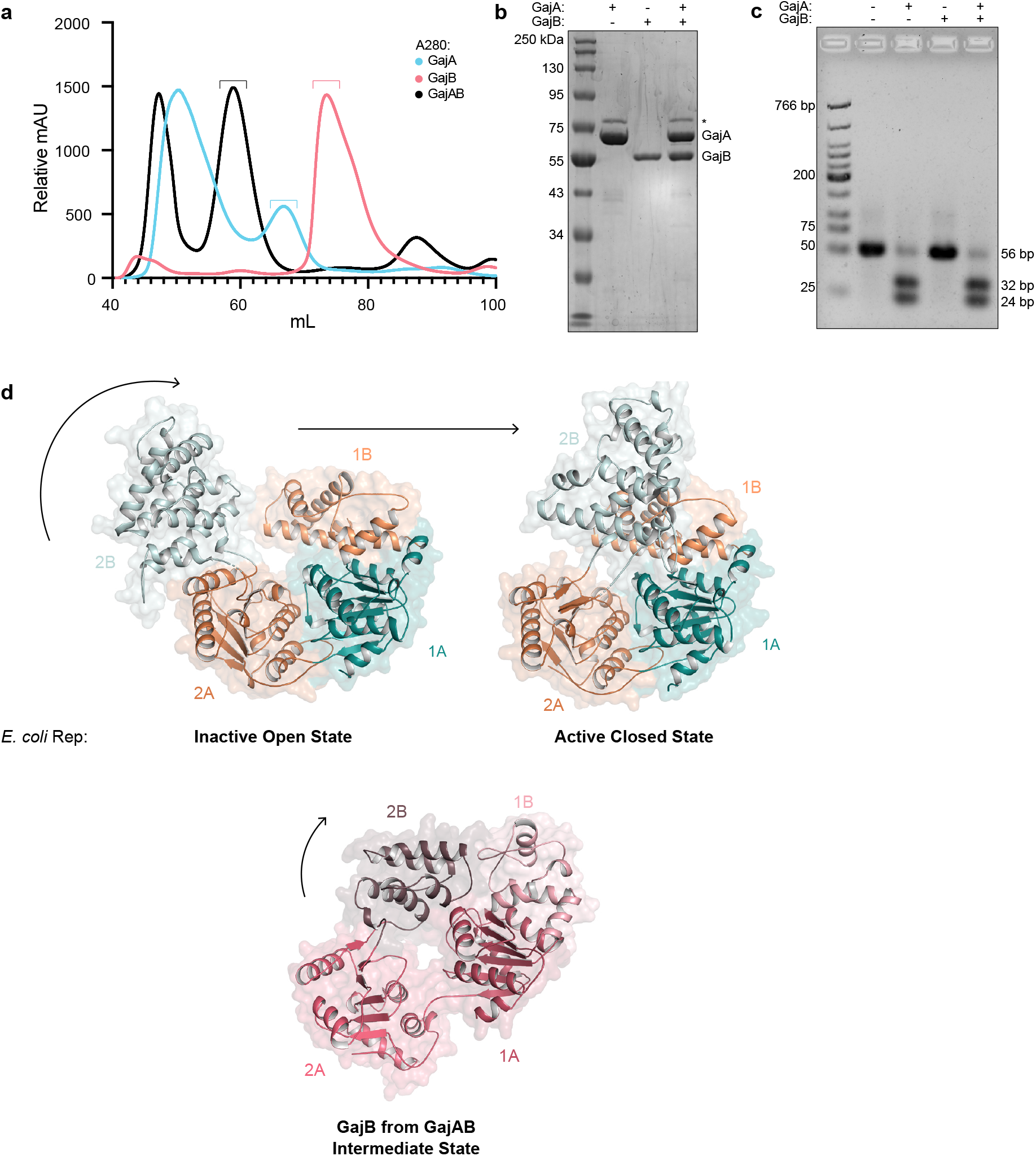
GajA and GajB form a supramolecular complex that cleaves phage lambda DNA *in vitro*. **a,** Size-exclusion chromatography (16/600 S200) analysis of recombinant *Bc*GajA and *Bc*GajB proteins, and the co-expressed *Bc*GajAB complex. Brackets indicate fractions collected for biochemical and structural analysis. **b,** SDS-PAGE analysis of purified GajA, GajB, and GajAB. Asterisk indicates minor contamination with the *E. coli* protein ArnA. **c,** Agarose gel analysis of the ability of GajA, GajB, and GajAB to cleave a 56-bp dsDNA demonstrates that GajA alone and the GajAB complex can cleave target DNA. **d,** Structural comparison of GajB and *Ec*Rep (PDB ID 1UAA)^19^ demonstrates the GajB 2B domain is rotated in a partially active intermediate position in the GajAB complex structure.

**Extended Data Figure 2.**
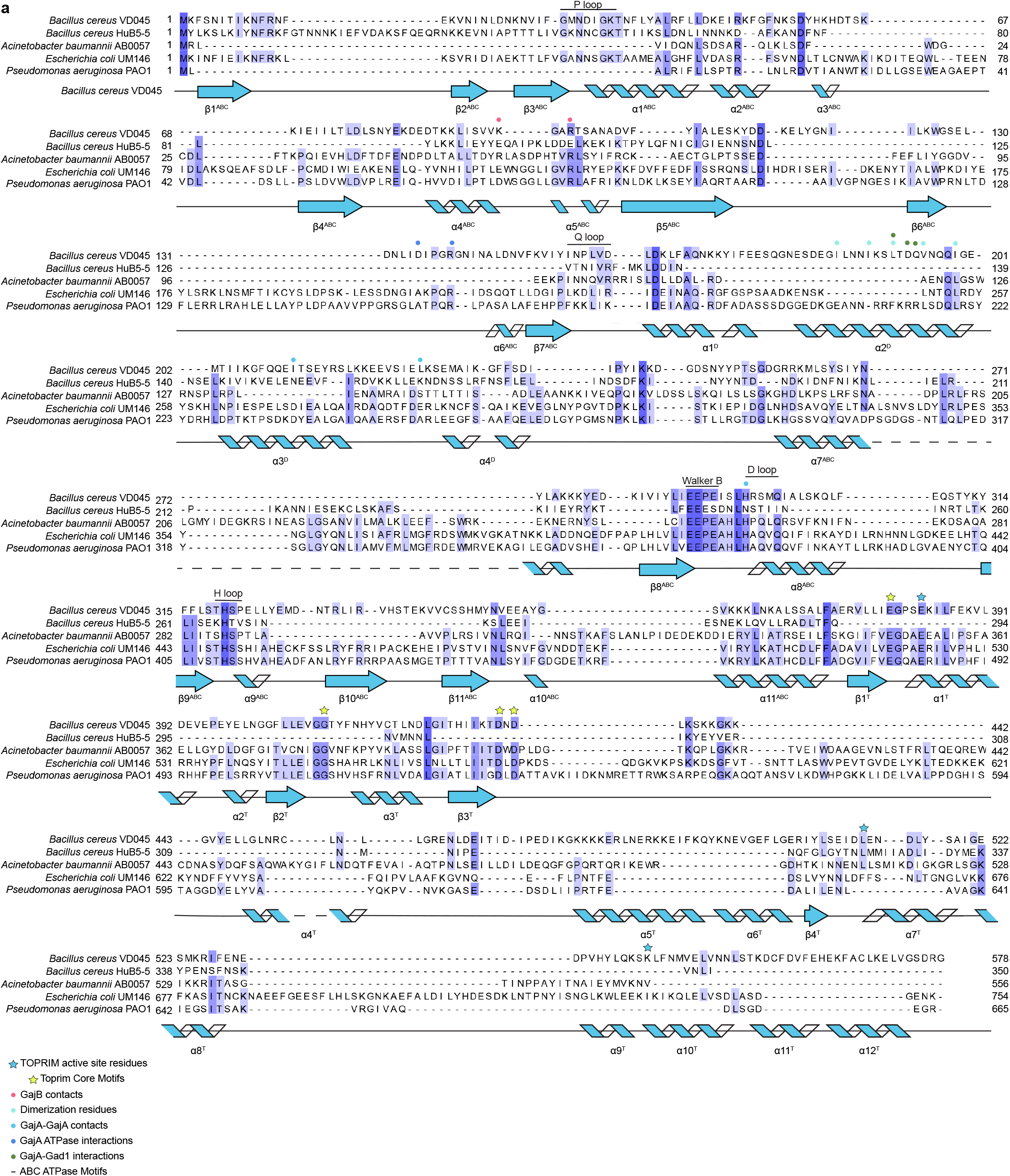
Structural characterization of GajA. **a,** Structure-guided alignment of GajA proteins from indicated bacteria colored according to amino acid conservation. The determined *Bacillus cereus* VD045 GajA secondary structure is displayed, and active-site and oligomerization interface residues are annotated according to the key below. Secondary structure abbreviations include ABC ATPase domain (ABC), dimerization domain (D), and Toprim domain (T).

**Extended Data Figure 3.**
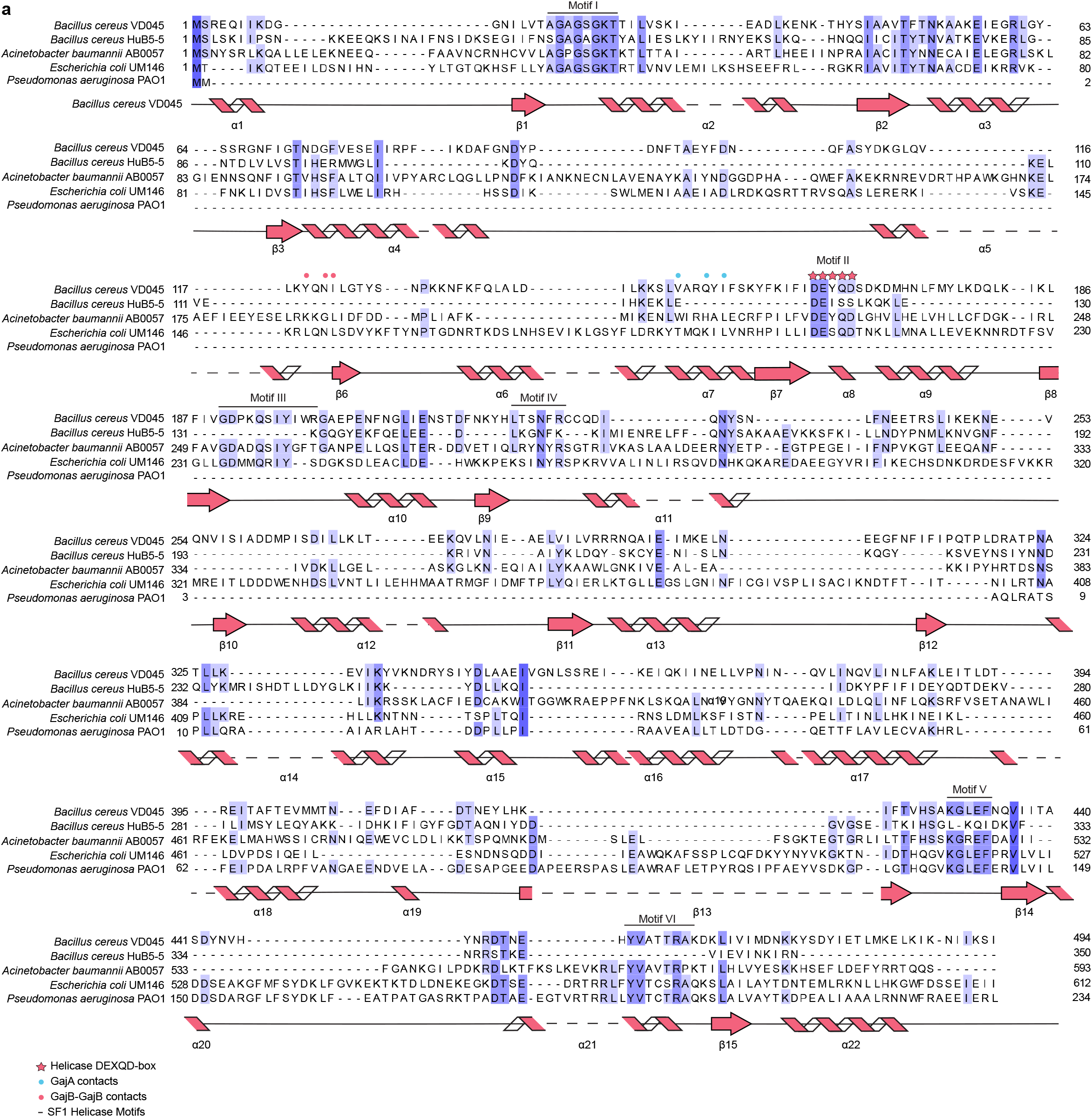
Structural characterization of GajB. **a,** Structure-guided alignment of GajB proteins from indicated bacteria colored according to amino acid conservation. The determined *Bacillus cereus* VD045 GajB secondary structure is displayed, and active-site and oligomerization interface residues are annotated according to the key below.

**Extended Data Figure 4.**
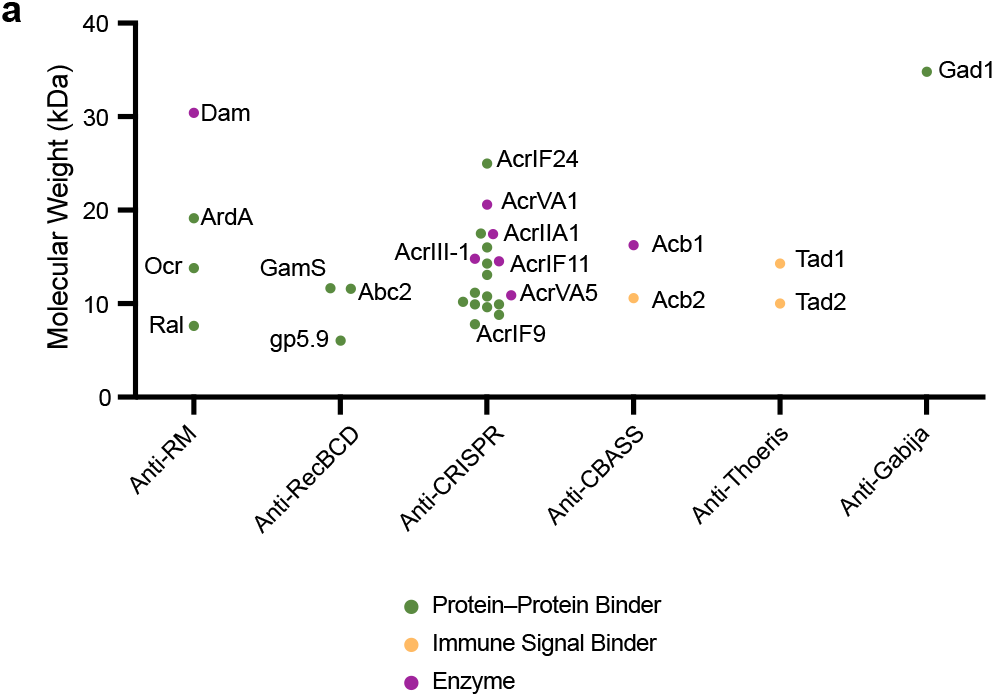
Size comparison of Gad1 to known phage immune evasion proteins. **a,** Analysis of known phage immune evasion proteins according to function and molecular weight demonstrates that Gad1 is atypically large for an evasion protein that functions through protein– protein interactions with a host anti-phage defense system. Phage immune evasion proteins are categorized and exhibited as colored dots colored according to the key below. Notable evasion proteins are indicated with text labels^23–27,40,47–50^.

**Extended Data Figure 5.**
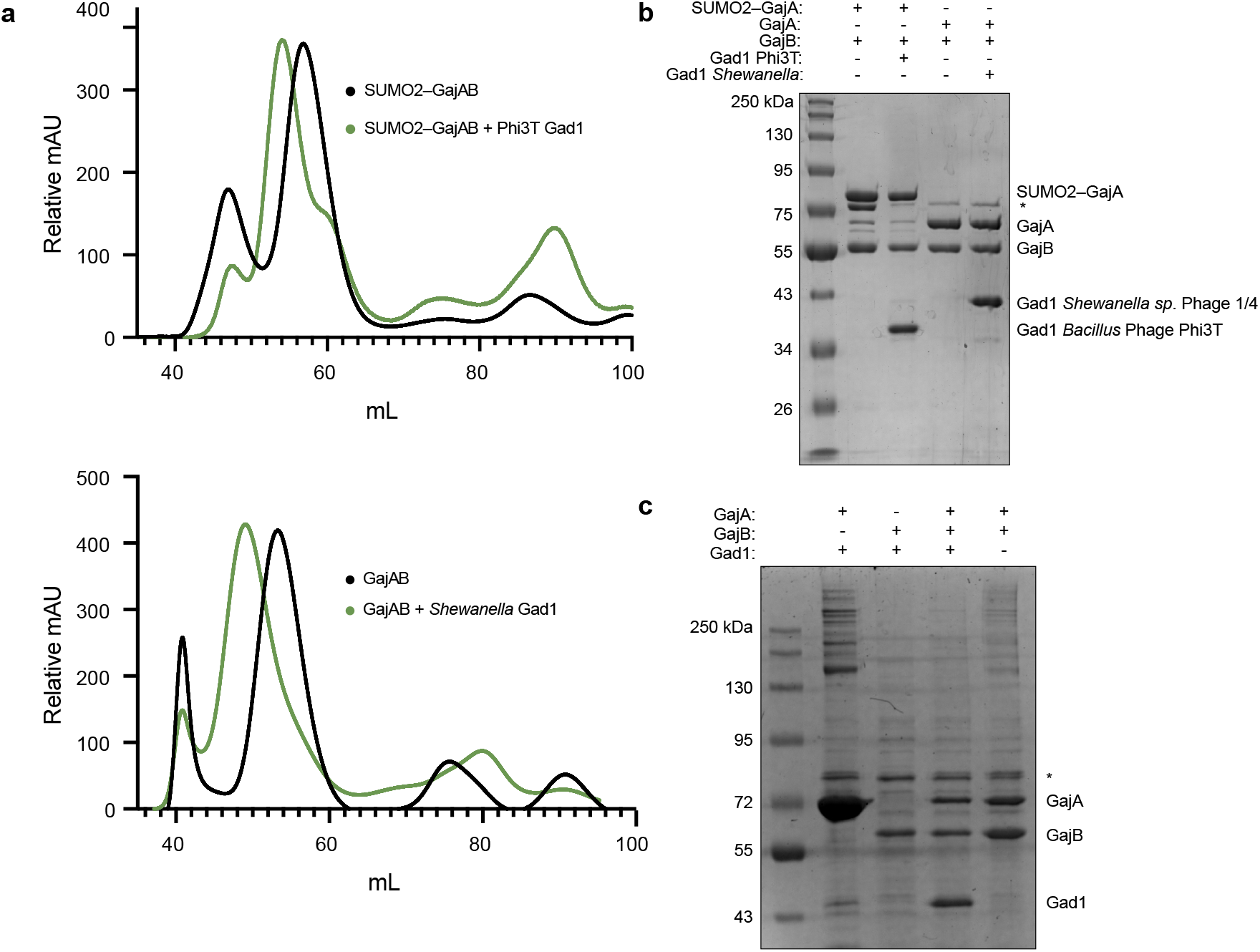
Biochemical characterization of Gad1 requirements for binding to the GajAB complex. **a,** Top, size-exclusion chromatography analysis (16/600 S200) of SUMO2-tagged *Bc*GajAB with or without phage Phi3T Gad1 used for cryo-EM structural studies. Bottom, size-exclusion chromatography analysis (16/600 S300) of *Bc*GajAB with or without *Shewanella* phage 1/4 Gad1 used for biochemical studies. *Shewanella* phage 1/4 Gad1 was used preferentially for biochemical studies due to less toxicity during *E. coli* expression. **b,** SDS-PAGE analysis of purified SUMO2- tagged GajAB, SUMO2-tagged GajAB in complex with phage Phi3T Gad1, untagged GajAB, and untagged GajB in complex with *Shewanella* phage 1/4 Gad1. **c,** SDS-PAGE analysis of Ni-NTA co-purified GajA, GajB, and GajAB with *Shewanella* phage 1/4 Gad1 indicates that Gad1 only binds the fully assembled GajAB complex. Asterisk indicates minor contamination with the *E. coli* protein ArnA.

**Extended Data Figure 6.**
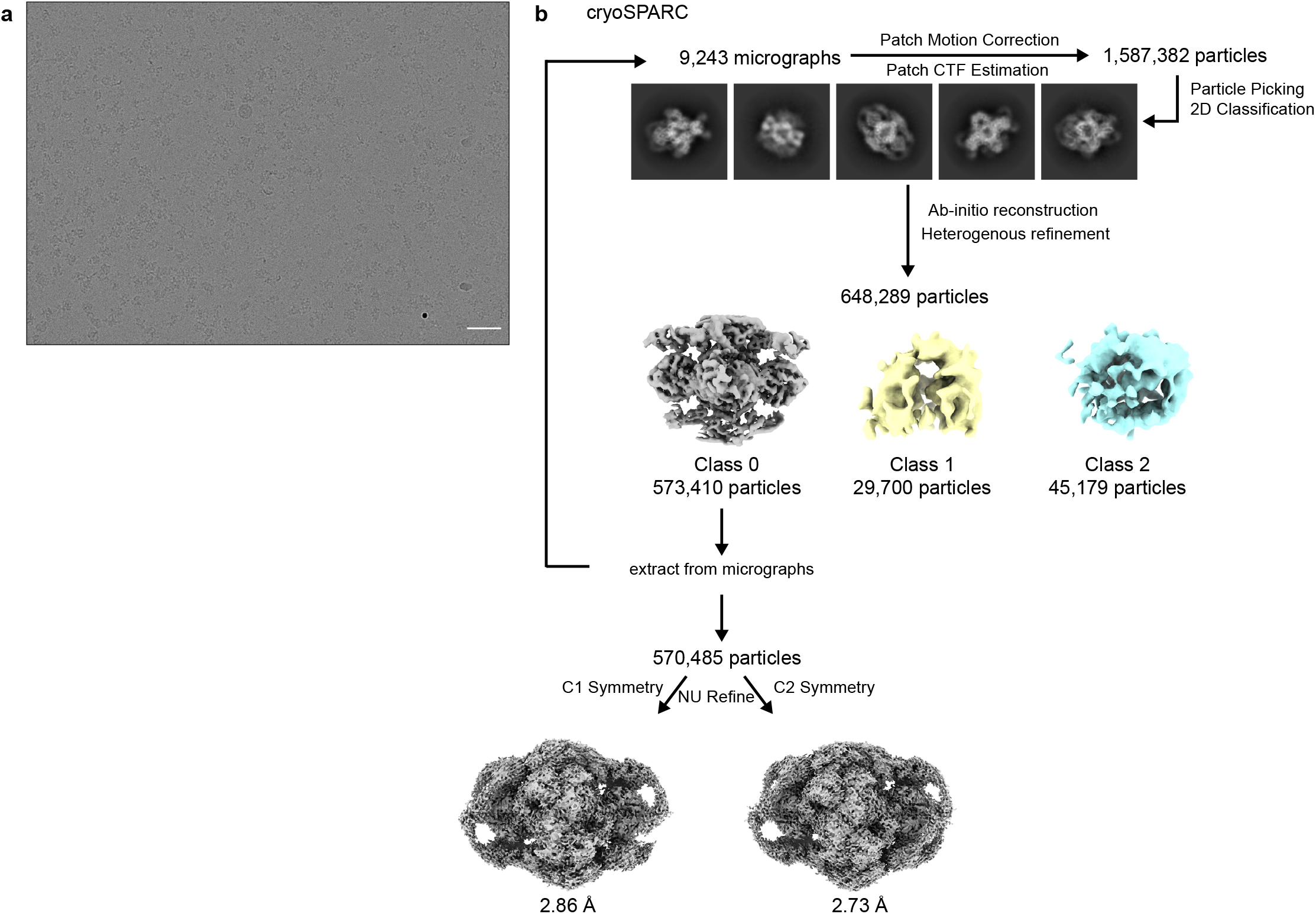
Cryo-EM data processing GajAB–Gad1 co-complex. **a,** Section of a representative electron micrograph (n = 9,243) of SUMO2–GajAB in complex with phage Phi3T Gad1. Scale bar is 50 nm. **b,** Data processing scheme used to generate the final 2.73 Å map.

**Extended Data Figure 7.**
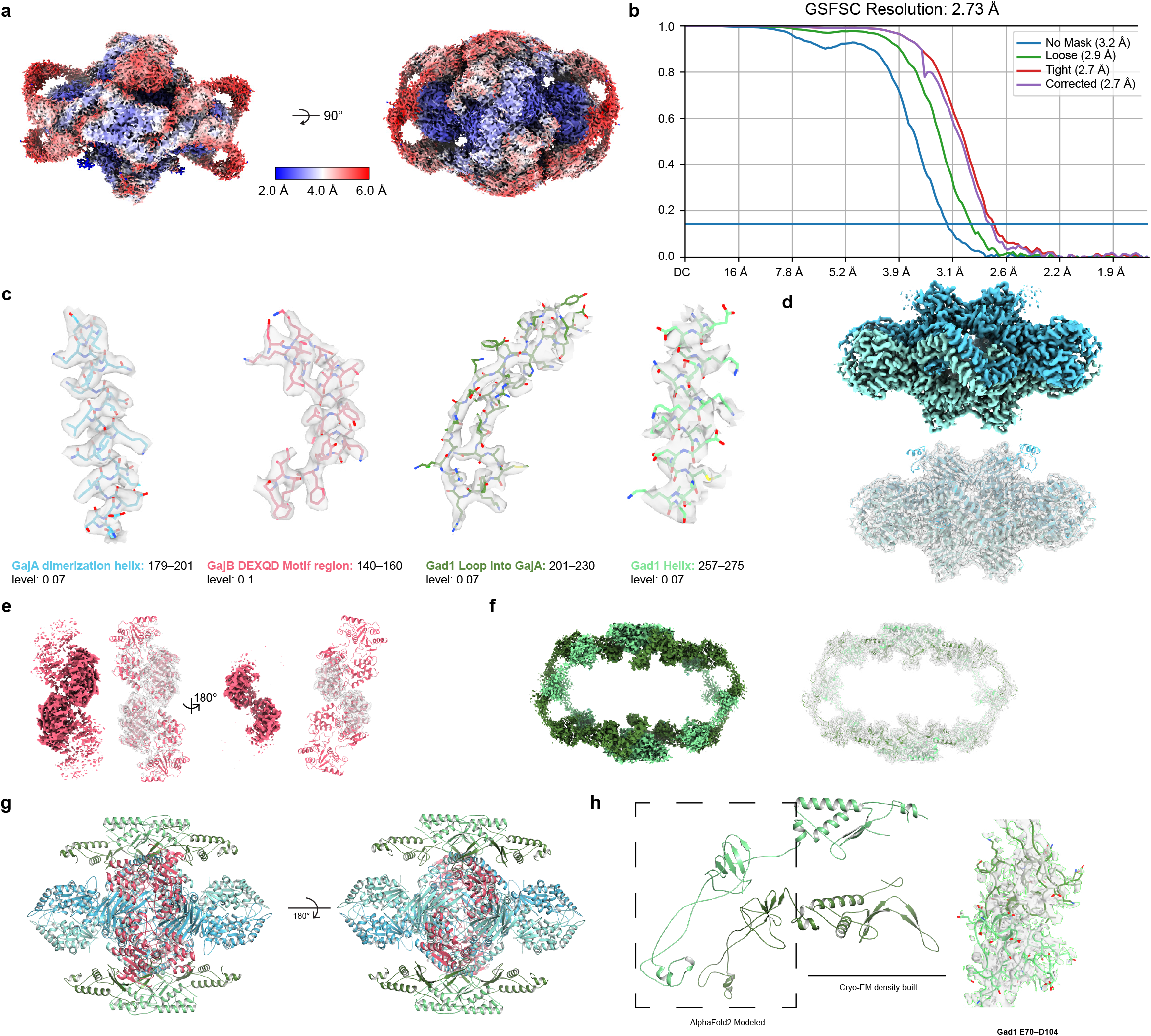
GajAB–Gad1 co-complex Cryo-EM map quality and model to map fitting. **a,** Reconstruction of the GajAB–Gad1 co-complex colored by local resolution. **b,** Fourier shell correlation (FSC) of the EM map. **c,** GajA, GajB, and Gad1 map to model fit for designated regions. **d,e,f,** Isolated GajA, GajB, Gad1 density maps with model fitting. **g,** GajAB–Gad1 model that was used for refining the cryo-EM map for Extended Data Table 2. **h,** Left, sections of Gad1 chains that were built *de novo* from the cryo-EM density and built using rigid-body placement of AlphaFold2 modeled residues. Right, cryo-EM density used to fit placement of Gad1 fist–fist domain contacts that complete protomer interactions.

**Extended Data Figure 8.**
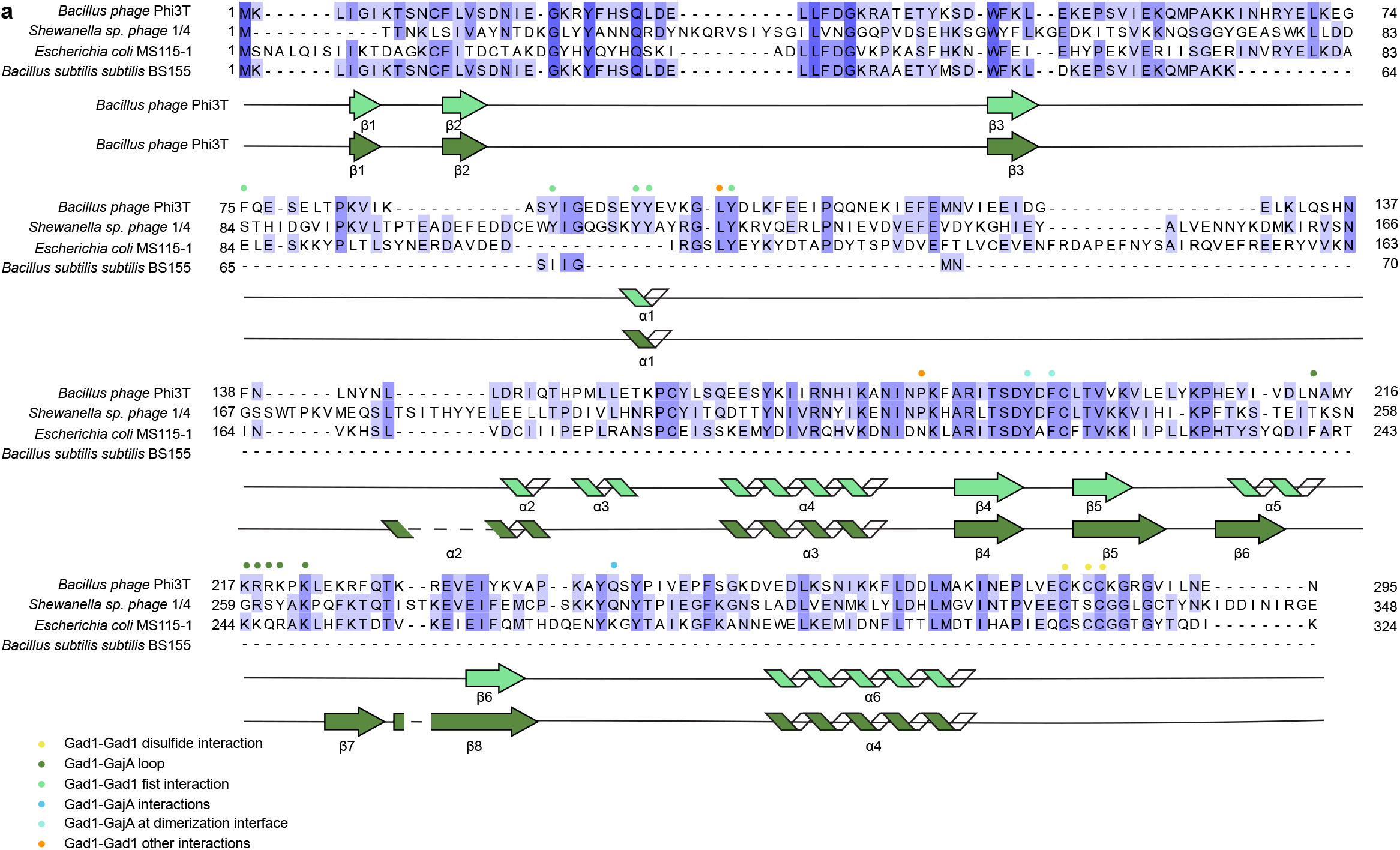
Structural characterization of Gad1. **a,** Structure-guided alignment of Gad1 proteins from indicated phage or prophage genomes colored according to amino acid conservation. The *Bacillus* phage Phi3T Gad1 secondary structure is displayed according to the two different conformations observed in the GajAB–Gad1 co-complex structure. Oligomerization interface residues are annotated according to the key below.

**Extended Data Figure 9.**
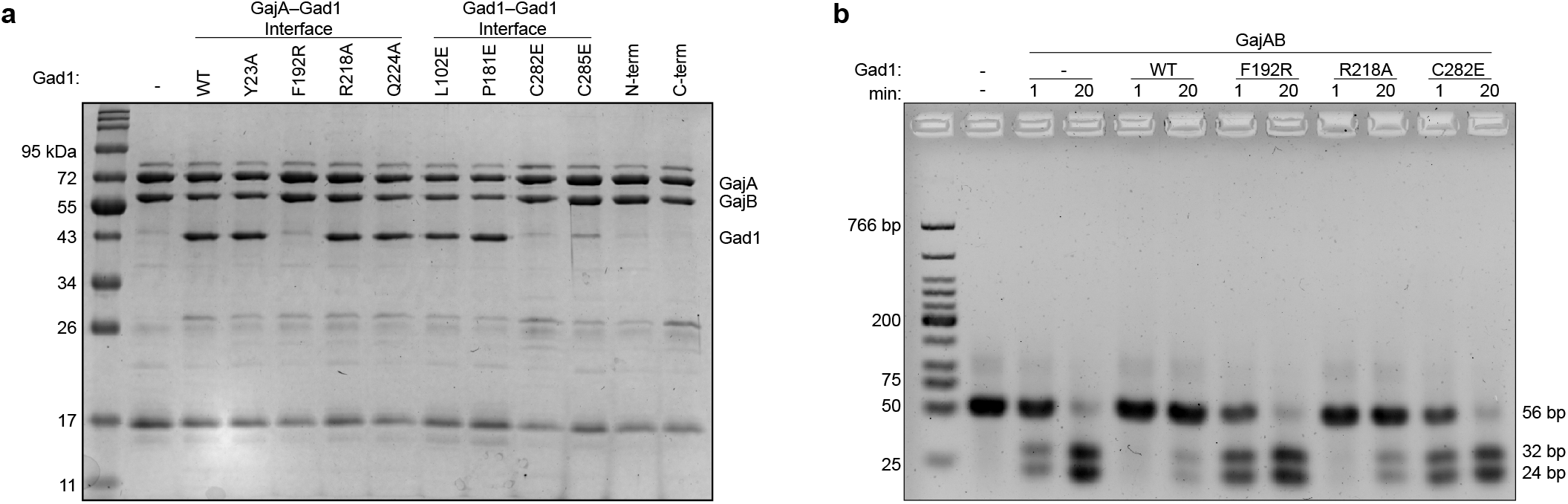
Biochemical characterization of Gad1 mutants that disrupt GajAB complex binding. **a,** SDS-PAGE analysis of the ability of *Shewanella* phage 1/4 Gad1 mutant proteins to interact with the GajAB complex. *Shewanella* phage 1/4 Gad1 mutant proteins were co-expressed with SUMO2-tagged GajAB (GajA-tagged) and co-purified by Ni-NTA pulldown. *Shewanella sp.* phage 1/4 Gad1 residues are numbered according to the Phi3T Gad1 structure. **b,** Agarose gel analysis of the ability of GajAB–Gad1 mutant complexes to cleave target 56-bp dsDNA after a minute and 20 minute incubation.

**Extended Data Figure 10.**
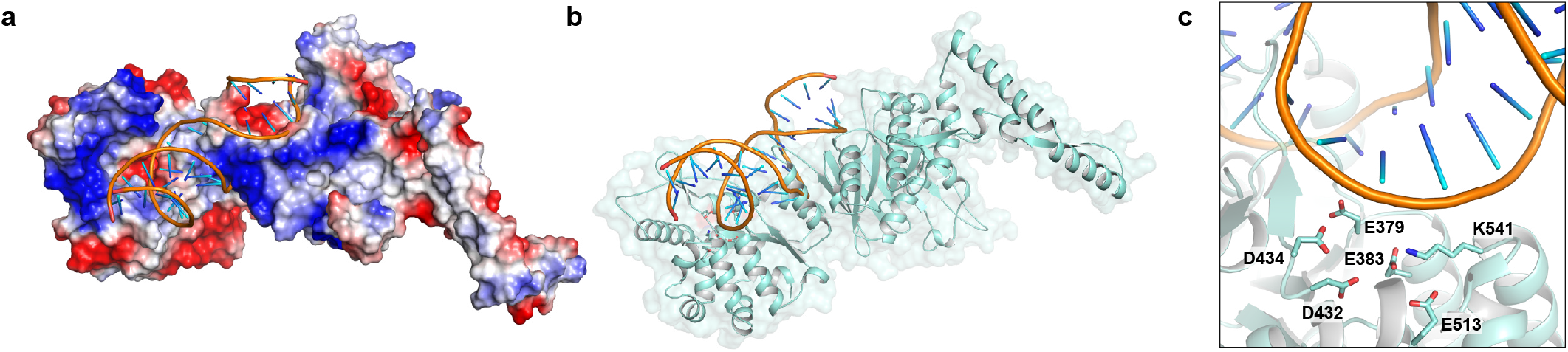
Modeling DNA-bound GajA. a,b, Isolated GajA protomer modeled with DNA bound to the Toprim domain shown with surface electrostatic potential (a) and in cartoon format (b). DNA modeling was performed using structural homology with the *E. coli* MutS–DNA complex (PDB ID 3K0S)^31^. **c,** Zoomed-in view of the GajA Toprim active site with modeled DNA.

**Extended Data Table 1.**
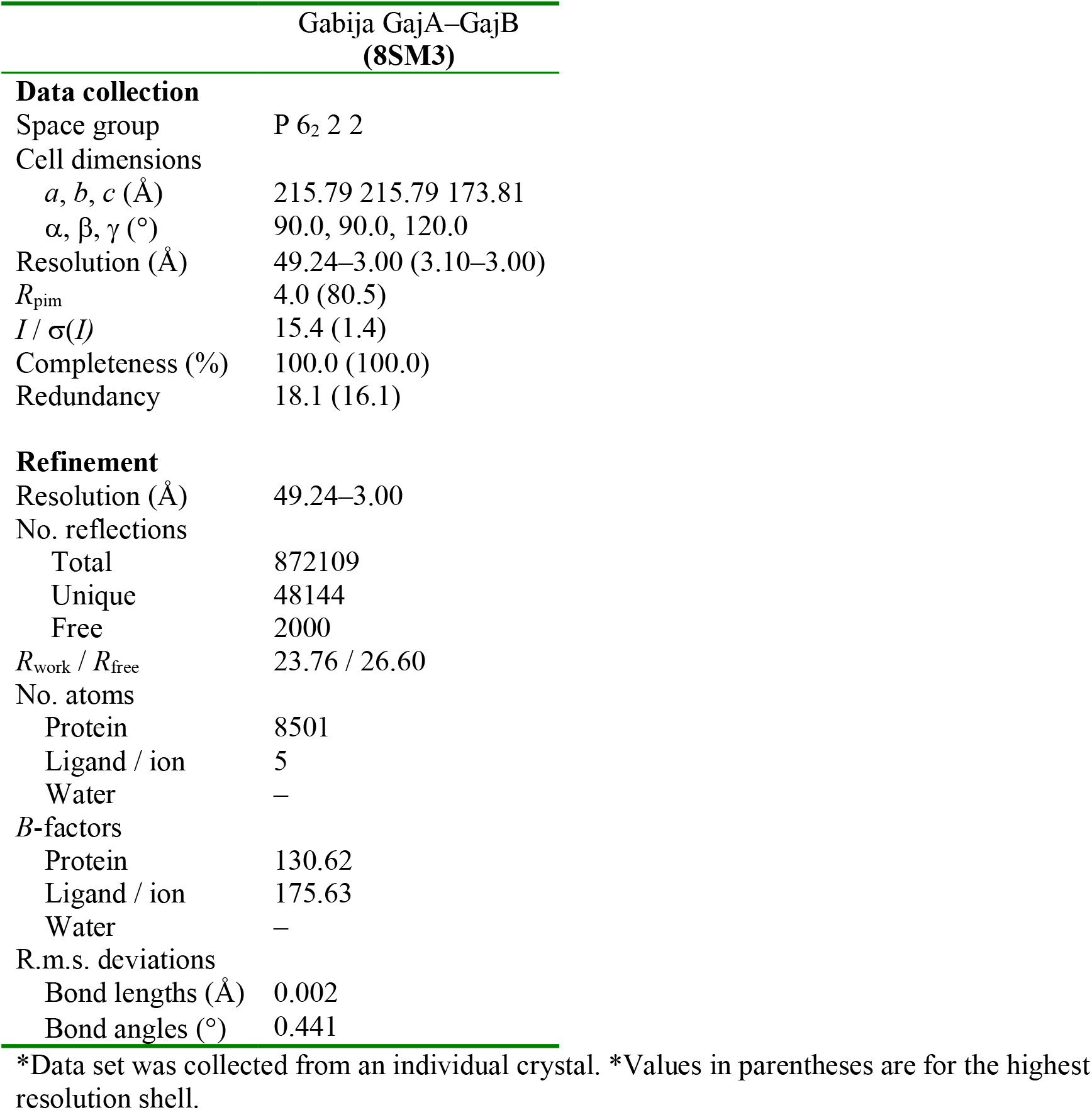
Summary of X-ray data collection, phasing and refinement statistics

**Extended Data Table 2.** Cryo-EM data collection, refinement and validation statistics

|  |  |
| --- | --- |
|  | GajAB-Gad1<br>co-complex<br>(EMD-xxxx)<br>(PDB xxxx) |
| **Data collection and processing** |  |
| Magnification | 37,000 |
| Voltage (kV) | 300 |
| Electron exposure (e-/Å<sup>2</sup>) | 41.1 |
| Defocus range (μm) | −0.8 to −1.9 |
| Pixel size (Å) | 0.3115 |
| Symmetry imposed | C2 |
| Initial particle images (no.) | 1,587,382 |
| Final particle images (no.) | 570,485 |
| Map resolution (Å) | 2.7 |
| FSC threshold | 0.143 |
| Map resolution range (Å) | 2.17–2.99 |
| **Refinement** |  |
| Initial model used (PDB code) |  |
| Model resolution (Å) | 2.71 |
| FSC threshold | 0.143 |
| Model resolution range (Å) | 2.71–2.73 |
| Map sharpening *B* factor (Å<sup>2</sup>) | −105.9 |
| Model composition |  |
| Non-hydrogen atoms | 35,487 |
| Protein residues | 4,338 |
| Ligands | 0 |
| *B* factors (Å<sup>2</sup>) |  |
| Proteins | 38.34 |
| R.m.s. deviations |  |
| Bond lengths (Å) | 0.003 |
| Bond angles (°) | 0.604 |
| Validation |  |
| MolProbity score | 1.95 |
| Clashscore | 9.55 |
| Poor rotamers (%) | 3.35 |
| Ramachandran plot |  |
| Favored (%) | 97.81 |
| Allowed (%) | 2.12 |
| Disallowed (%) | 0.07 |

